# Empirical test of crab-clam predator-prey model predictions: storm-driven phase shift to a low-density steady state

**DOI:** 10.1101/224097

**Authors:** Cassandra N. Glaspie, Rochelle D. Seitz, Romuald N. Lipcius

## Abstract

A dynamic systems approach can predict steady states in predator-prey interactions, but there are very few empirical tests of predictions from predator-prey models. Here, we examine the empirical evidence for the low-density steady state predicted by a Lotka-Volterra model of a crab-clam predator-prey system using data from long-term monitoring, a field survey, and a field experiment. We show that Tropical Storm Agnes in 1972 likely resulted in a phase shift to a low-density state for the soft-shell clam *Mya arenaria*, which was once a biomass dominant in Chesapeake Bay. This storm altered predator-prey dynamics between *M. arenaria* and the blue crab *Callinectes sapidus*, shifting from a system controlled from the bottom-up by prey resources, to a system controlled from the top-down by predation pressure on bivalves. Predator-prey models with these two species alone were capable of reproducing observations of clam densities and mortality rates, consistent with the idea that *C. sapidus* are a major driver of *M. arenaria* population dynamics. Over 40 y post-storm, *M. arenaria* densities hover near a low-density steady state predicted from the predator-prey model. Relatively simple models can predict phase shifts and identify alternative stable states, as shown by agreement between model predictions and field data in this system. The preponderance of multispecies interactions exhibiting nonlinear dynamics indicates that this may be a general phenomenon.

## 1. INTRODUCTION

Predators play a key role in ecosystem stability and function by consuming dominant competitors (Lubchenco & Gaines 1981, Boudreau & Worm 2012). Predators can also destabilize ecosystems or collapse food webs if they become too abundant, or if their prey do not have natural defenses against predation. Generally, predators and their prey have evolved over time to coexist. Prey have anti-predator behaviors or morphological adaptations to avoid being eaten (Bibby et al. 2007, Whitlow 2010). Similarly, predators have adaptations or behaviors that help them to forage optimally and take advantage of prey when they are available (Meire & Ervynck 1986, Rindone & Eggleston 2011).

One of the ways the balance between predator and prey adaptations manifests in nature is through density-dependent predation. Predators can exhibit a numerical response to prey densities by increasing reproduction rates due to an overabundance of prey (demographic response) or by gathering in areas with relatively high densities of prey (aggregative response) (Holling 1959). An individual predator may also adjust its predation rate to prey density through a ‘functional response’ (changes in a predator’s consumption rate in response to prey density). Density-dependent mechanisms tend to stabilize prey population dynamics (Royama 1992, Turchin 2003) and can maintain population viability when a population is reduced to low levels (Cushing 1975).

Certain characteristics of a predator-prey system can help predict which functional response will be observed. A linear relationship between consumption rate and prey density (type I functional response) is expected for organisms that do not actively search for prey, such as filter feeders. Most vertebrate and invertebrate predators exhibit a hyperbolic functional response that increases to an asymptote due to limits associated with prey handling, ingestion, and metabolism (type II functional response) (Hassell et al. 1977). Predators that feed upon cryptic or otherwise hard-to-find prey exhibit a sigmoidal functional response, where consumption rates increase slowly at low prey densities (type III functional response) (Holling 1959). Prey that avoid predators can achieve a low-density refuge; thus, the functional response can explain the distribution of prey items, and it can be used to predict the persistence of prey species at low densities (Eggleston et al. 1992).

There are many mathematical models that can be used to predict predator-prey dynamics (Briggs & Hoopes 2004). These models contain nonlinear functions describing the density-dependent interactions between predator and prey. Due to these nonlinearities, model behavior often includes shifts to alternative stable states (Drake & Griffen 2010). These states may include extinction of one or both species, or coexistence steady states where both predator and prey are able to coexist at densities predicted by the model. Multiple stable coexistence states are possible in fairly simple predator-prey functions (Mumby et al. 2007, Kramer & Drake 2010).

A dynamic systems approach can predict coexistence states in natural and managed systems. Steady states analysis been used to develop optimum harvesting and conservation strategies for forest and rangeland systems (Hritonenko et al. 2013, Bauch et al. 2016). The theory surrounding steady states and bifurcations is well developed, but there are very few empirical tests. To date, empirical tests of coexistence steady states in predator-prey systems have involved populations of small organisms such as unicellular organisms, yeast, small crustaceans, or bacteria (Luckinbill 1973, van den Ende 1973, Drake & Griffen 2010). Identifying or establishing coexisting predator-prey systems for use in testing model predictions has proven difficult, especially for macro-organisms. Of the few empirical tests of steady states that focus on macro-organisms, none examines interactions between natural populations of predators and prey (Kramer & Drake 2010, Jiang et al. 2018, McNickle & Evans 2018). Here, we show that a severe storm resulted in a phase shift to a low-density state for the soft-shell clam *Mya arenaria*, which was once a biomass dominant in Chesapeake Bay, in the face of predation by the blue crab *Callinectes sapidus*. We examine the empirical evidence for the low-density steady state predicted by a Lotka-Volterra model of this predator-prey system using data from long-term monitoring, a field survey, and a field experiment.

### 1.1. History of predator-prey system

Tropical Storm Agnes, which reached and remained in the Chesapeake Bay watershed 21-23 June 1972, has long been suspected of resulting in long-term changes for the Bay (Orth & Moore 1983). Tropical Storm Agnes was a “100-year storm” that caused sustained, extremely low salinities (Figure 1) and increased sedimentation throughout Chesapeake Bay (Schubel 1976, Schubel et al. 1976). This storm has been blamed for the loss of seagrass in certain areas of Chesapeake Bay (Orth & Moore 1983), high mortality rates and recruitment failure in oysters *Crassostrea virginica* (Haven et al. 1976), and declines in abundance of the soft-shell clam *Mya arenaria*, which suffered a mass mortality after the storm (Cory & Redding 1976).

**Figure 1.**
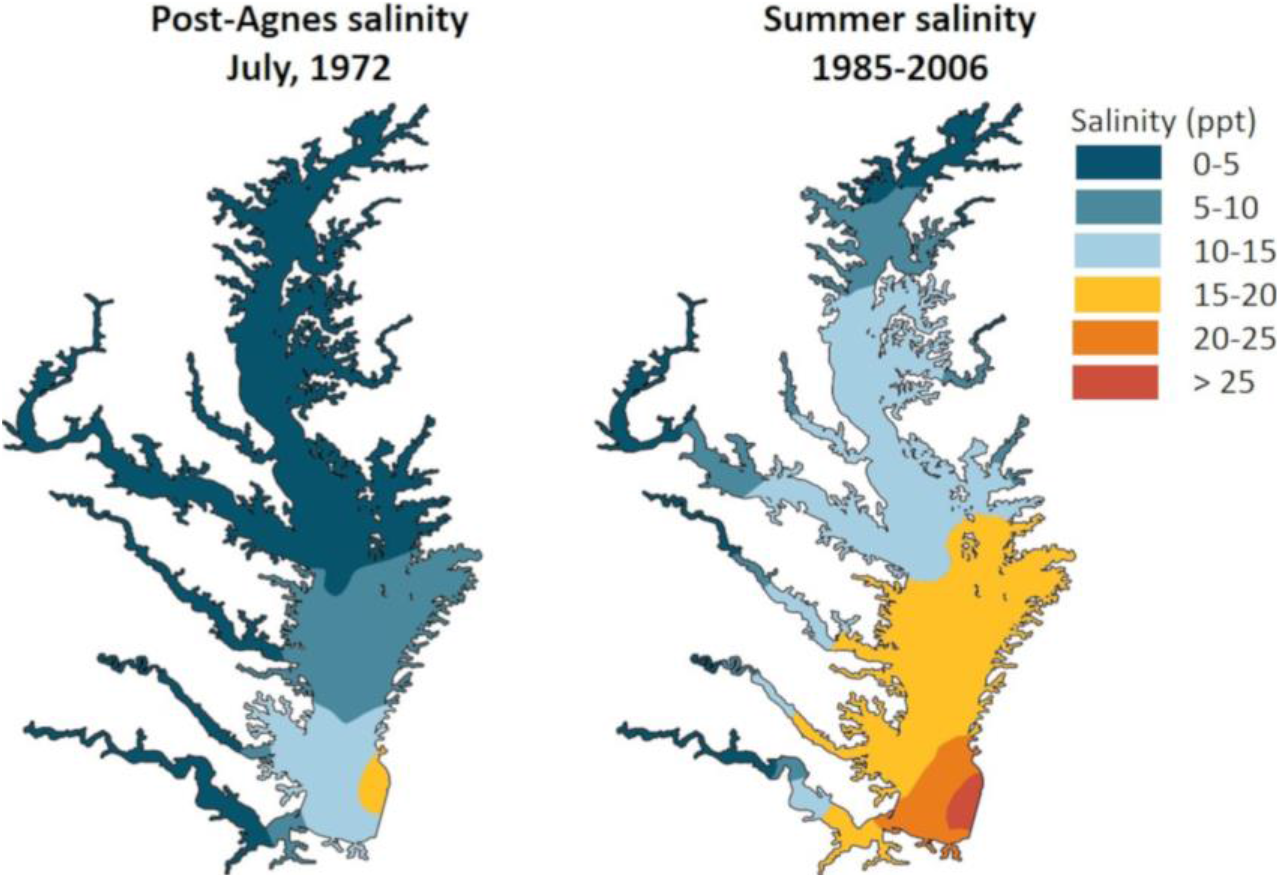
Pre- and post- storm salinity profiles. Salinity profiles for post-Agnes (left) and average summer (right) conditions. Post-Agnes salinity was measured over the period June 29 – July 3, 1972 (Schubel et al. 1976). The summer salinity profile (right) is average surface salinity for 1985-2006 (Weinberg 2008).

*Mya arenaria* was abundant enough to support a major commercial fishery throughout Chesapeake Bay prior to 1972 (Haven 1970). When the population declined abruptly after Tropical Storm Agnes, the fishery never recovered in lower Chesapeake Bay (Virginia) (Glaspie et al. 2018). Attempts to revive a commercial fishery in Virginia waters were never realized after the storm’s passage. The commercial fishery for soft-shell clams in the Maryland portion of the Bay is characterized by variable and low harvest (Dungan et al. 2002); the fishery declined by 89% after the storm and has been near collapse since (NMFS).

The failure of *M. arenaria* to recover from storm-related declines has been attributed to predation, habitat loss, disease, rising temperatures, and overfishing (Glaspie et al. 2018). The Virginia and Maryland portions of Chesapeake Bay have different habitats, disease dynamics, climates, and fishing pressure; therefore, these factors are unlikely to explain the inability of *M. arenaria* to recover from low density in both regions (Dungan et al. 2002, Glaspie et al. 2018). More recently, disease has been blamed for an added minor decline in *M. arenaria* (Dungan et al. 2002); however, there is no evidence that disease prevalence or intensity are correlated with *M. arenaria* density (Glaspie et al. 2018).

Experimental evidence suggests that on a local scale, interactions between *M. arenaria* and their major predator, the blue crab *C. sapidus* (Meise & Stehlik 2003), are capable of keeping clams at low densities (Lipcius & Hines 1986, Seitz et al. 2001). *Mya arenaria* burrow deeply in sediments, and when clams are at low densities, crabs are unable to detect their presence (Lipcius & Hines 1986). The result is a low-density refuge for *M. arenaria*, driven by disproportionately low predation, which is characteristic of a sigmoidal functional response (Lipcius & Hines 1986, Seitz et al. 2001). Given this evidence regarding a potential mechanism for the decline in *M. arenaria* and maintenance of the population at low density, this study examines the empirical evidence for a low-density steady state in this predator-prey system, and the impact of Tropical Storm Agnes on basin-scale population dynamics of *M. arenaria*.

## 2. MATERIALS & METHODS

Changepoint analysis of time series was conducted using R statistical software on *M. arenaria* landings (NMFS 2017) and log-transformed average adult female *C. sapidus* abundance (VIMS trawl survey) in the Chesapeake Bay from 1958-1992, with an AIC penalty and using the segment neighbor algorithm (Auger & Lawrence 1989). This time period was chosen for analysis because it begins when *M. arenaria* landings data first became available and ends before the slow decline in landings in the early 1990s due to fisheries collapse.

Predator-prey ordinary differential equation (ODE) models were modified with a type III functional response:

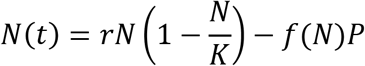

where N is the density of prey, P is the density of predators, r is the intrinsic per capita growth rate, K is the carrying capacity, and *f*(*N*) takes the form of a sigmoidal functional response:

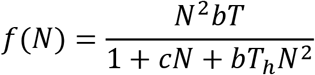

where T is the time available for foraging, T_h_ is handling time, and b and c are components of the attack rate in a sigmoidal functional response (Hassell et al. 1977).

Models were parameterized using data from the literature as follows: P = 0.06 m^−2^ (MD DNR), r = 1.75 y^−1^ (Brousseau 1978), K = 200 m^−2^ (Abraham & Dillon 1986), T = 1 y, Th = 0.0015 y (Lipcius & Hines 1986), b = 26.30 y^−1^ (Lipcius & Hines 1986), and c = 0.14 (Lipcius & Hines 1986). Analytical solutions of steady states were calculated using Matlab statistical software. Stability of each coexistence steady state was determined by examining the sign of eigenvalues.

To examine mortality rates, we solved the equation for number consumed:

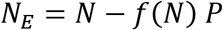

where N_E_ = the number of clams eaten calculated for a period of 8 d (0.022 y) at an initial density of N = 48 m^−2^ to match the field predation experiments (Glaspie & Seitz 2018). We then calculated mortality as:

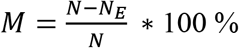

where M = percent mortality. Density of predators P was allowed to vary to achieve M = 76.3% (Glaspie & Seitz 2018), and the resultant predator density that achieved observed mortality rates of juvenile *M. arenaria* was compared to published *C. sapidus* densities for Chesapeake Bay. Data and R code files are available in the Knowledge Network for Biocomplexity (KNB) repository (Glaspie 2019).

## 3. RESULTS

Tropical Storm Agnes in 1972 resulted in a phase shift for *M. arenaria*, which was maintained at low abundance likely due to predation by the blue crab *C. sapidus*. An abrupt shift in clam abundance was identified in 1972, the year of Tropical Storm Agnes (Figure 2). Before the storm, crab abundance was positively correlated with clam abundance at a lag of 1 y (r = 0.66, p = 0.01), indicating that each year, clams were prey for juvenile crabs that recruited to the fishery at one year of age (Figure 3a). After the storm, clam abundance was negatively correlated with crab abundance with a lag of 1 y (r = −0.48, p = 0.04), indicating that each year, crabs were consuming juvenile clams that would have recruited to the fishery a year later (Figure 3b). This is consistent with a phase shift from a system controlled from the bottom-up by prey resources, to a system controlled from the top-down by predation pressure on bivalves.

**Figure 2.**
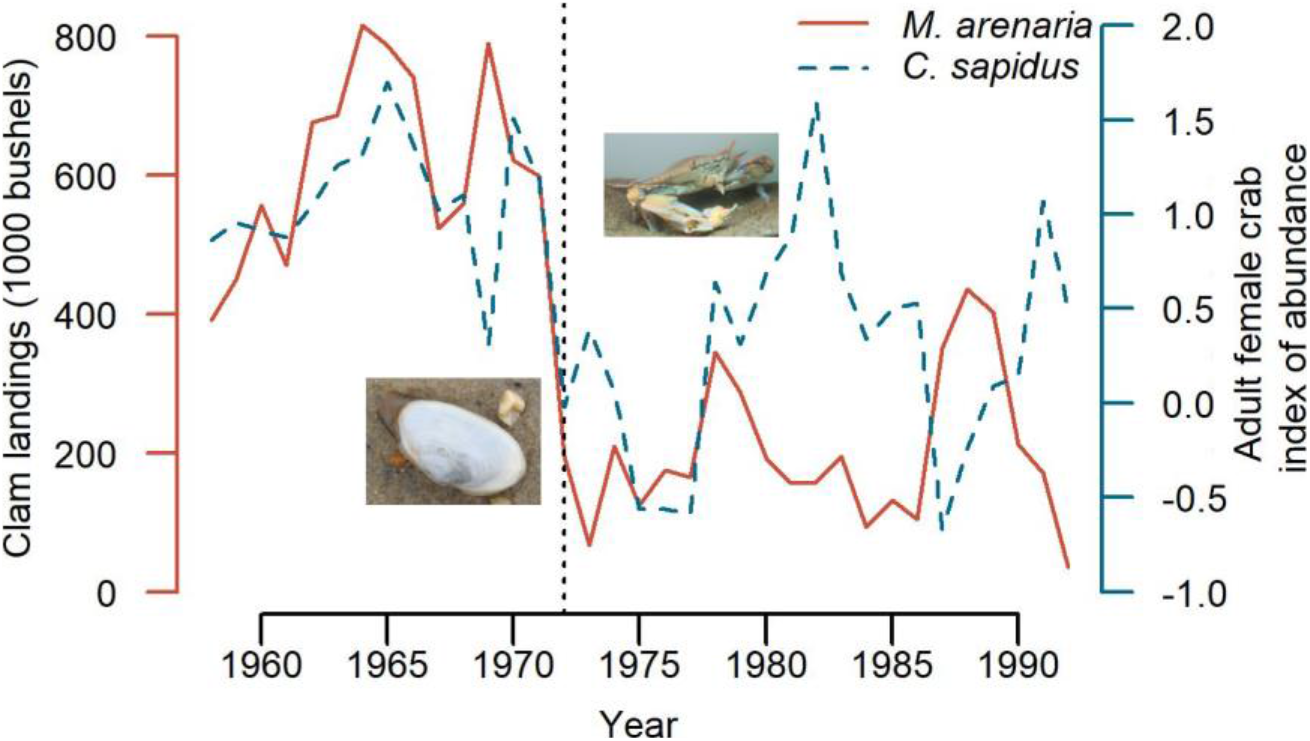
Predator-prey time series. Time series for soft-shell clam *Mya arenaria* landings (red) and adult female blue crab *Callinectes sapidus* index of abundance (blue). Blue crab data are log-transformed average female abundance per tow (VIMS trawl survey). *Mya arenaria* data are fisheries landings (1000 bushels) (NMFS Annual Commercial Landing Statistics). Vertical dashed line represents Tropical Storm Agnes (1972), and the location of the changepoint from time series analysis. Photo credits: C.N. Glaspie (clam), R.N. Lipcius (crab).

**Figure 3.**
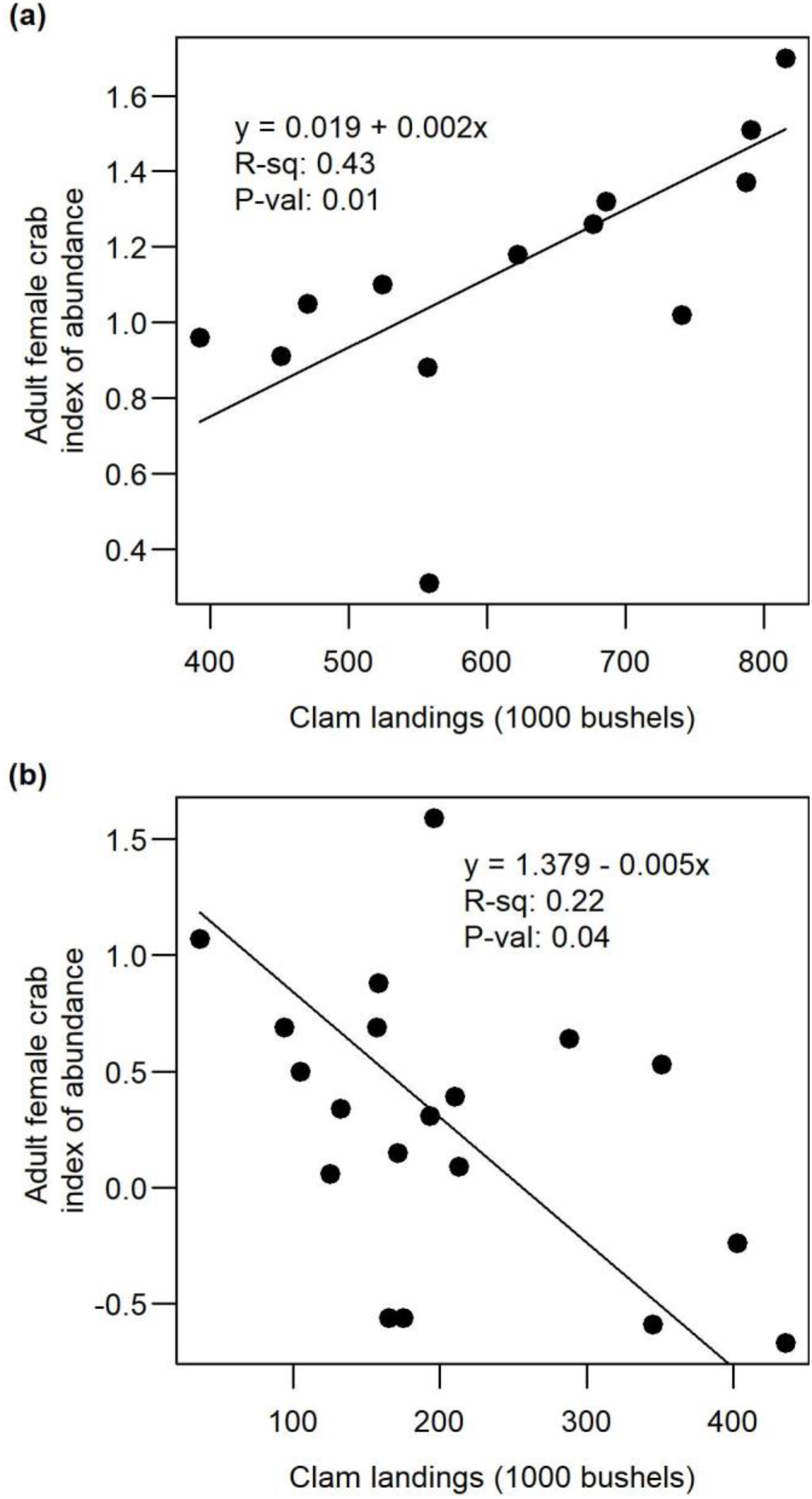
Pre- and post- storm relationship between crab index of abundance (log-transformed average female abundance per tow, VIMS trawl survey) and clam landings (fisheries-dependent data). (a) Before the storm, crab abundance was positively correlated with clam abundance (1 y lag). (b) After the storm, clam abundance was negatively correlated with crab abundance with (1 y lag).

Predator-prey modeling confirmed the presence of high-density (near carrying capacity at 173.99 clams m^−2^) and low-density (at 1.41 clams m^−2^) steady states separated by an unstable steady state at 20.93 clams m^−2^ (Figure 4). We propose that *M. arenaria* existed in Chesapeake Bay at high densities until perturbed past the unstable steady state in 1972 by Tropical Storm Agnes. Thereafter, it was able to persist at low density due to the low-density refuge from blue crab predation (Lipcius & Hines 1986, Seitz et al. 2001), rather than collapsing to local extinction. Unfortunately, *M. arenaria* is unlikely to rebound to high abundance without a beneficial disturbance, such as a considerable recruitment episode or substantial reduction in predation pressure, which propels it above the unstable steady state (Figure 4) and concurrently allows it to overcome the exacerbated disease burden.

**Figure 4.**
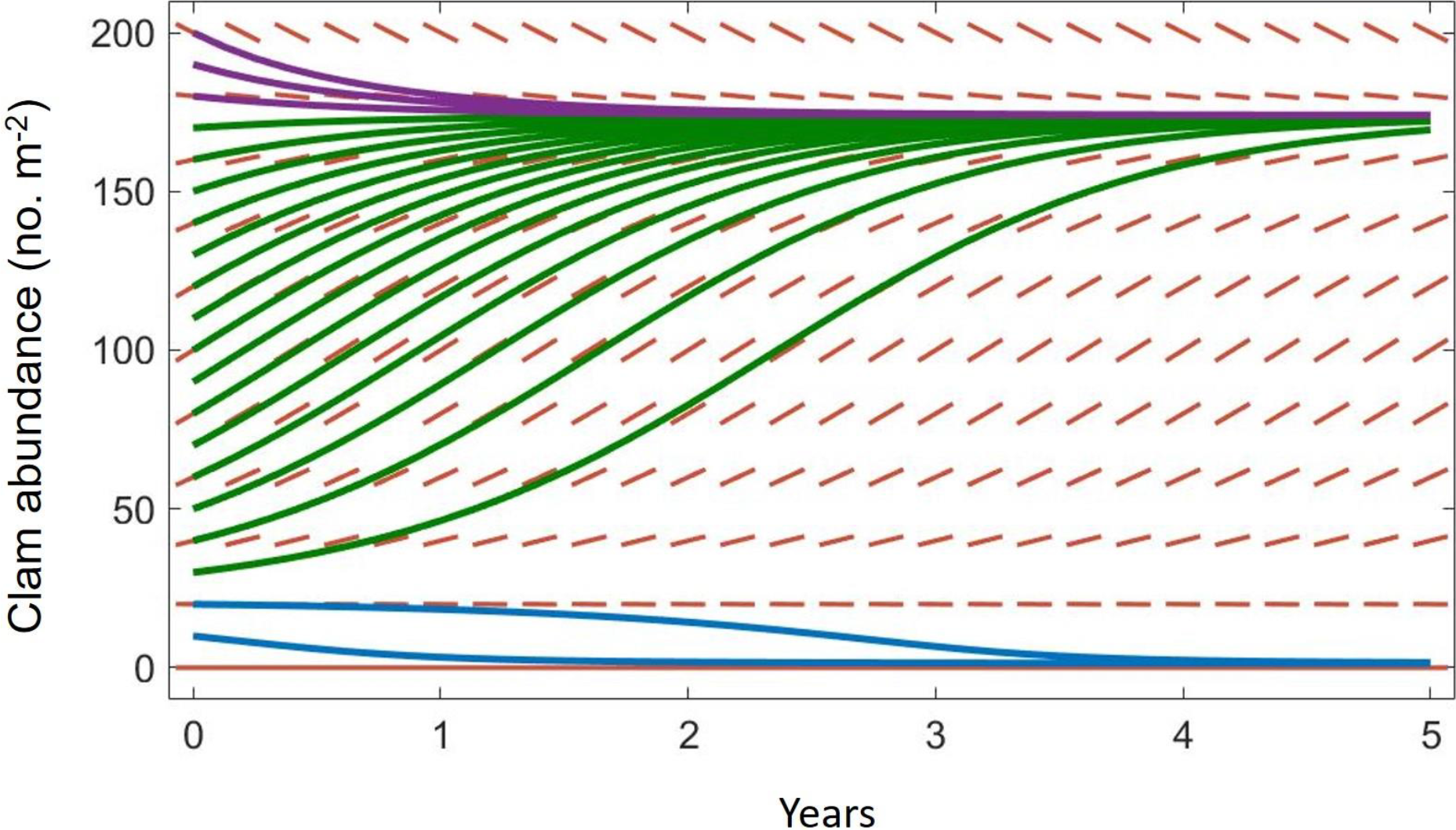
Slope field diagrams for predator-prey models. Trajectories of *Mya arenaria* density either approach a high-density stable steady state at carrying capacity (green and purple lines) or a low-density stable steady state at 1.41 clams m^−2^. Trajectories diverge from an unstable steady state at 20.93 clams m^−2^.

Predator-prey models with these two species alone were capable of reproducing observations of clam densities and mortality rates, consistent with the idea that blue crabs are a major driver of *M. arenaria* population dynamics (Lipcius & Hines 1986, Seitz et al. 2001, Meise & Stehlik 2003). The low-density steady state predicted by the predator-prey model is similar to observed densities of *M. arenaria* in Chesapeake Bay. *Mya arenaria* persist in the lower Chesapeake Bay at average densities of 0.4 – 1.73 m^−2^ (95% CI), despite episodes of high recruitment (Glaspie et al. 2018). In the field, juvenile *M. arenaria* exposed to predators suffered 76.3% higher mortality as compared to caged individuals (Glaspie & Seitz 2018). Predator exclusion treatments confirmed that blue crabs were responsible for most of the mortality of juvenile *M. arenaria* (Glaspie & Seitz 2018), and mortality rates observed in the field were comparable to mortality rates predicted by the model for blue crab density 4.8 m^−2^, which is a typical density for juvenile crabs in the summer months in Chesapeake Bay (Ralph & Lipcius 2014).

## 4. DISCUSSION

The observations, theory, and mechanistic basis indicate that *M. arenaria* was subjected to a storm-driven phase shift to low density, which has been maintained by blue crab predation in Chesapeake Bay. Extreme weather events are costly, and they are likely to become even more common with predicted increases in the intensity and frequency of extreme events due to anthropogenic climate change (Settele et al. 2014). When examining the cost of extreme weather, ecological impacts are rarely considered, even though the impacts of such events on the ecosystem may be severe (Thomson et al. 2012). Evidence for storm-driven phase shifts in coral reefs (Mumby et al. 2007), kelp ecosystems (Byrnes et al. 2011), and now soft-sediment communities (current study) suggests that management of ecosystems should include an examination of nonlinear interactions and the potential for phase shifts.

To our knowledge, this is one of a handful of empirical tests of predator-prey theory as a predictive tool in natural systems. More evidence is needed to fully evaluate the predictive potential of steady-state analysis involving predator-prey models, and the value of a dynamics systems approach to ecosystem modeling efforts. However, the approach described here can be adapted to a variety of predator-prey systems with available estimates of life history parameters, population density, and distribution. The approach may also be expanded to include more than two species, or encompass additional complexity, such as metabolic functions.

The concepts encapsulated in the crab-clam predator-prey system, including population self-limitation, consumer-resource oscillations, and the functional response are widespread in nature. These concepts may be considered “laws” of population ecology (Turchin 2001). Each concept listed above introduces nonlinear dynamics into population models, especially those with multiple interacting species. Given the preponderance of multispecies interactions exhibiting nonlinear dynamics, multiple steady states may be a general ecological phenomenon.

## Supporting information

WebTable 1

## Acknowledgments

We gratefully acknowledge assistance by the students and staff of the Community Ecology and Marine Conservation Biology labs at the Virginia Institute of Marine Science. CNG performed the analysis and wrote the manuscript. CNG, RDS, and RNL contributed substantially to study conception and design, interpretation of the data, and manuscript revisions. This material is based upon work supported by the National Oceanic and Atmospheric Administration grant number NA11NMF4570218; the Environmental Protection Agency EPA STAR Fellowship under grant number FP91767501; and the National Science Foundation Administration GK-12 program under grant number DGE-0840804. This paper is contribution number XXXX from the Virginia Institute of Marine Science, College of William & Mary.

## Notes

#### Summary of Updates

This version of the manuscript has been revised for submission to Marine Ecology Progress Series.

https://knb.ecoinformatics.org/#view/doi:10.5063/F19S1PB9

## References

Abraham BJ, Dillon PL (1986) Species profiles: Life histories and environmental requirements of coastal fishes and invertebrates (Mid-Atlantic)- softshell clam. U.S. Fish and Wildlife Service Biological Report 82(11.68)

Auger IE, Lawrence CE (1989) Algorithms for the optimal identification of segment neighborhoods. Bull Math Biol 51:39–54

Bauch CT, Sigdel R, Pharaon J, Anand M (2016) Early warning signals of regime shifts in coupled human–environment systems. Proc Natl Acad Sci 113:14560–14567

Bibby R, Cleall-Harding P, Rundle S, Widdicombe S, Spicer J (2007) Ocean acidification disrupts induced defences in the intertidal gastropod *Littorina littorea*. Biol Lett 3:699–701

Boudreau SA, Worm B (2012) Ecological role of large benthic decapods in marine ecosystems: A review. Mar Ecol Prog Ser 469:195–213

Briggs CJ, Hoopes MF (2004) Stabilizing effects in spatial parasitoid-host and predator-prey models: A review. Theor Popul Biol 65:299–315

Brousseau DJ (1978) Population dynamics of the soft-shell clam *Mya arenaria*. Mar Biol 50:63–71

Byrnes JE, Reed DC, Cardinale BJ, Cavanaugh KC, Holbrook SJ, Schmitt RJ (2011) Climate-driven increases in storm frequency simplify kelp forest food webs. Glob Chang Biol 17:2513–2524

Cory RL, Redding MJ (1976) Mortalities caused by Tropical Storm Agnes to clams and oysters in the Rhode River area of Chesapeake Bay. In: Anderson AM (ed) The Effects of Tropical Storm Agnes on the Chesapeake Bay Estuarine System. The Johns Hopkins University Press, Baltimore, MD, p 478–487

Cushing D (1975) Marine Ecology and Fisheries. Cambridge University Press, NY

Drake JM, Griffen BD (2010) Early warning signals of extinction in deteriorating environments. Nature 467:456–459

Dungan CF, Hamilton RM, Hudson KL, McCollough CB, Reece KS (2002) Two epizootic diseases in Chesapeake Bay commercial clams, *Mya arenaria* and *Tagelus plebeius*. Dis Aquat Organ 50:67–78

Eggleston DB, Lipcius RN, Hines AH (1992) Density-dependent predation by blue crabs upon infaunal clam species with contrasting distribution and abundance patterns. Mar Ecol Prog Ser 85:55–68

Ende P van den (1973) Predator-prey interactions in continuous culture. Science 181:562–564

Glaspie CN (2019) Blue crab *Callinectes sapidus* and soft-shell clam *Mya arenaria* index of abundance time series 1958-1992. Knowl Netw Biocomplexity:doi: 10.5063/F19S1PB9

Glaspie CN, Seitz RD (2018) Habitat complexity and benthic predator-prey interactions in Chesapeake Bay. PLoS One 13:e0205162

Glaspie CN, Seitz RD, Ogburn MB, Dungan CF, Hines AH (2018) Impacts of habitat, predators, recruitment, and disease on soft-shell clams *Mya arenaria* and stout razor clams *Tagelus plebeius* in Chesapeake Bay. Mar Ecol Prog Ser 603:117–133

Hassell MP, Lawton JH, Beddington JR (1977) Sigmoid functional responses by invertebrate predators and parasitoids. J Anim Ecol 46:249–262

Haven DS (1970) A study of the hard and soft clam resources of Virginia: Annual contract report for the period 1 July 1969 through 30 June 1970. Gloucester Point, VA

Haven DS, Hargis WJ, Loesch JG, Whitcomb JP (1976) The effect of Tropical Storm Agnes on oysters, hard clams, soft clams, and oyster drills in Virginia. In: Anderson AM (ed) The Effects of Tropical Storm Agnes on the Chesapeake Bay Estuarine System. The Johns Hopkins University Press, Baltimore, MD, p 488–508

Holling C (1959) The components of predation as revealed by a study of small mammal predation of the European pine sawfly. Can Entomol 91:293–320

Hritonenko N, Yatsenko Y, Goetz RU, Xabadia A (2013) Optimal harvesting in forestry: Steady-state analysis and climate change impact. J Biol Dyn 7:41–58

Jiang J, Huang Z-G, Seager TP, Lin W, Grebogi C, Hastings A, Lai Y-C (2018) Predicting tipping points in mutualistic networks through dimension reduction. Proc Natl Acad Sci:201714958

Kramer AM, Drake JM (2010) Experimental demonstration of population extinction due to a predator-driven Allee effect. J Anim Ecol 79:633–639

Lipcius RN, Hines AH (1986) Variable functional responses of a marine predator in dissimilar homogenous microhabitats. Ecology 67:1361–1371

Lubchenco J, Gaines SD (1981) A unified approach to marine plant-herbivore interactions. I. Populations and communities. Annu Rev Ecol Syst 12:405–437

Luckinbill L (1973) Coexistence in laboratory populations of *Paramecium aurelia* and its predator *Didinium nasutum*. Ecology 54:1320–1327

McNickle GG, Evans WD (2018) Toleration games: Compensatory growth by plants in response to enemy attack is an evolutionarily stable strategy. AoB Plants 10:1–14

Md DNR 2017 Blue Crab Winter Dredge Survey.

Meire PM, Ervynck A (1986) Are oystercatchers (*Haemoptopus ostralegus*) selecting the most profitable mussels (*Mytilus edulis*)? Anim Behav 34:1427–1435

Meise CJ, Stehlik LL (2003) Habitat use, temporal abundance variability, and diet of blue crabs from a New Jersey estuarine system. Estuaries 26:731–745

Mumby PJ, Hastings A, Edwards HJ (2007) Thresholds and the resilience of Caribbean coral reefs. Nature 450:98–101

NMFS Annual Commercial Landing Statistics. https://www.st.nmfs.noaa.gov/commercial-fisheries/commercial-landings (accessed 10-27-2017)

NMFS (2017) Fisheries Economics of the United States 2015. Washington, D.C.

Orth RJ, Moore KA (1983) Chesapeake Bay: An unprecedented decline in submerged aquatic vegetation. Science (80−) 222:51–53

Ralph GM, Lipcius RN (2014) Critical habitats and stock assessment: Age-specific bias in the Chesapeake Bay blue crab population survey. Trans Am Fish Soc 143:889–898

Rindone RR, Eggleston DB (2011) Predator-prey dynamics between recently established stone crabs (*Menippe* spp.) and oyster prey (*Crassostrea virginica*). J Exp Mar Bio Ecol 407:216–225

Royama T (1992) Analytical Population Dynamics. Chapman & Hall, London, UK

Schubel JR (1976) Effects of Agnes on the suspended sediment on the Chesapeake Bay and contiguous shelf waters. In: Schubel JR (ed) The Effects of Tropical Storm Agnes on the Chesapeake Bay Estuarine System. The Johns Hopkins University Press, Baltimore, MD, p 179–200

Schubel JR, Carter HH, Cronin WB (1976) Effects of Agnes on the distribution of salinity along the main axis of the Bay and in contiguous shelf waters. In: Ruzecki EP (ed) The Effects of Tropical Storm Agnes on the Chesapeake Bay Estuarine System. The Johns Hopkins University Press, Baltimore, MD, p 33–65

Seitz RD, Lipcius RN, Hines AH, Eggleston DB (2001) Denity-dependent predation, habitat variations, and the persistence of marine bivalve prey. Ecology 82:2435–2451

Settele J, Scholes R, Betts R, Bunn S, Leadley P, Nepstad D, Overpeck JT, Taboada MA (2014) Terrestrial and inland water systems. In: Field CB, Barros VR, Dokken DJ, Mach KJ, Mastrandrea MD, Bilir TE, Chatterjee M, Ebi KL, Estrada YO, Genova RC, Girma B, Kissel ES, Levy AN, MacCracken S, Mastrandrea PR, White LL (eds) Climate Change 2014: Impacts, Adaptation, and Vulnerability. Part A: Global and Sectoral Aspects. Contribution of Working Group II to the Fifth Assessment Report of the Intergovernmental Panel on Climate Change. Cambridge University Press, New York, NY, p 271–359

Thomson JR, Bond NR, Cunningham SC, Metzeling L, Reich P, Thompson RM, Nally R Mac (2012) The influences of climatic variation and vegetation on stream biota: Lessons from the Big Dry in southeastern Australia. Glob Chang Biol 18:1582–1596

Turchin P (2001) Does population ecology have general laws? Oikos 94:17–26

Turchin P (2003) Complex Population Dynamics. Princeton University Press, Princeton, New Jersey

Weinberg H (2008) Map: Chesapeake Bay Mean Surface Salinity - Summer (1985-2006). http://www.chesapeakebay.net/maps/map/https://www.chesapeakebay.net/what/maps/chesapeake_bay_mean_surface_salinity_summer_1985_2006 (accessed 10-29-2017)

Whitlow WL (2010) Changes in survivorship, behavior, and morphology in native soft-shell clams induced by invasive green crab predators. Mar Ecol 31:418–430

