## Supplementary material for "Empirical test of crab-clam predator-prey model predictions: storm-driven phase shift to a low-density steady state": WebTable 1

| Parameter | Value | Units | Reference |
| --- | --- | --- | --- |
| P | 0.06 | m^-2^ | MD DNR |
| r | 1.75 | y^-1^ | Brousseau 1978 |
| K | 200 | m^-2^ | Abraham and Dillon 1986 |
| T | 1 | y |  |
| T_h_ | 0.0015 | y | Lipcius and Hines 1986 |
| b | 26.30 | y^-1^ | Lipcius and Hines 1986 |
| c | 0.14 | unitless | Lipcius and Hines 1986 |

WebReferences:

Abraham BJ and Dillon PL. 1986. Species profiles: Life histories and environmental requirements of coastal fishes and invertebrates (Mid-Atlantic)- softshell clam. U.S. Fish and Wildlife Service Biological Report 82(11.68).

Brousseau DJ. 1978. Population dynamics of the soft-shell clam Mya arenaria. Mar Biol 50: 63–71.

Lipcius RN and Hines AH. 1986. Variable functional responses of a marine predator in dissimilar homogenous microhabitats. Ecology 67: 1361–71.

MD DNR. 2017 Blue Crab Winter Dredge Surveyhttp://dnr.maryland.gov/fisheries/Pages/blue-crab/dredge.aspx. Viewed 29 Oct 2017.
